# Osmophoresis – a possible mechanism for vesicle trafficking in tip-growing cells

**DOI:** 10.1101/029157

**Authors:** Andrei Lipchinsky

## Abstract

A mechanism for polarized transport of vesicles by means of osmotic propulsions is proposed and substantiated for tip-growing cells. An analysis is presented which shows that in pollen tubes the gradient of cytosolic water potential can drive vesicle movement either in the anterograde or retrograde direction, depending on the vesicle position, its radius and the phase of growth oscillation. The importance of transcellular water flow for cytoskeletal dynamics and cell motility is highlighted.

---

This is an author-created, un-copyedited version of an article published in Physical Biology 12 (2015) 066012. The Version of Record is available online at http://dx.doi.org/10.1088/1478-3975/12/6/066012

Tip growth is an extreme form of polarized cell growth characterized by cell surface extension restricted to a small area at the cell tip. Such a growth pattern is responsible for producing filamentous cells with enormous length-to-width ratio: fungal hyphae, algal rhizoids, moss and fern protonemata, root hairs, pollen tubes and some other highly polarized structures (Rounds and Bezanilla, 2013; Domozych et al., 2013; Sanati Nezhad and Geitmann, 2015). Tip-growing cells can sustain prodigious rates of elongation, up to tens of micrometers per minute, as in the case of pollen tubes and fungal hyphae (Jauh and Lord, 1995; Lew, 2011). Since cell surface expansion occurs in a restricted area (< 80 μm^2^), the high growth rates imply that plasma membrane and cell wall constituents are expelled from the growing apical dome into non-growing cell regions within a few minutes after their deposition. This constitutive drift is coupled to concerted vesicular delivery of materials into the growing apex from older cell regions. The vesicular transport along non-growing cell regions up to the flanks of the apical dome is ensured by motor proteins that convey cargoes on well-defined longitudinal cytoskeletal tracks (Campanoni and Blatt, 2007; Domozych et al., 2013; Rounds and Bezanilla, 2013). In the cell apex, however, cytoskeletal network is sparse, and tip-focused gradient of Ca^2+^ promotes actin turnover and can inactivate myosin, making the involvement of motor proteins in the terminal stage of vesicle trafficking doubtful (Kroeger et al., 2009; Chebli et al., 2013; Cai et al., 2015; Hepler and Winship, 2015). Therefore, it has been suggested that in the cell apex the motion of vesicles is governed by a combination of advection by the cytosol and diffusion (Kroeger et al., 2009). However, such an assumption perhaps overlooks osmotic heterogeneity of the apical cytoplasm as a potential factor involved in vesicular transport. The objective of this study is to highlight the role of cytosolic osmotic gradient in vesicle dynamics in pollen tubes as model tip-growing cells that are well characterized with respect to osmotically important ion currents.

A striking pattern of localized Ca^2+^, H^+^, K^+^ and Cl^−^ fluxes in pollen tubes has been described and shown to be implicated in the regulation of growth processes and vesicle motility (e.g. Lazzaro et al., 2005; Lovy-Wheeler et al., 2006; Breygina et al., 2009a; Tavares et al., 2011). The magnitude of Cl^−^ fluxes greatly exceeds the magnitude of the other ion fluxes, indicating that Cl^−^ dynamics is the major driver of osmotic gradients within pollen tubes. Strong oscillatory Cl^−^ efflux occurs at the tube apex, while steady Cl^−^ influx takes place along the tube shank starting at ~12 μm apart from the tip (Fig. 1A). The peak magnitude of oscillatory Cl^−^ efflux varies in time and in a cell-specific manner from 1 up to 60 nmol·cm^−2^·s^−1^ (Zonia et al., 2002; see also Kunkel et al. 2006; Moreno et al., 2007; Michard et al., 2009; Tavares et al., 2011; Gutermuth et al., 2013). Based upon these flux values and using Fick’s law, the concentration gradients of cytosolic Cl^−^ can be tentatively estimated to be 10^1^ − 6·10^5^ mol·m^−4^ in the tube tip and 3·10^4^ − 2·10^6^ mol·m^−4^ in the region spanning from the flanks of the apical dome to the beginning of the zone where Cl^−^ influx is started (Appendix 1). This gradient can be a key driver of vesicle dynamics, as explained below.

**Fig. 1.**
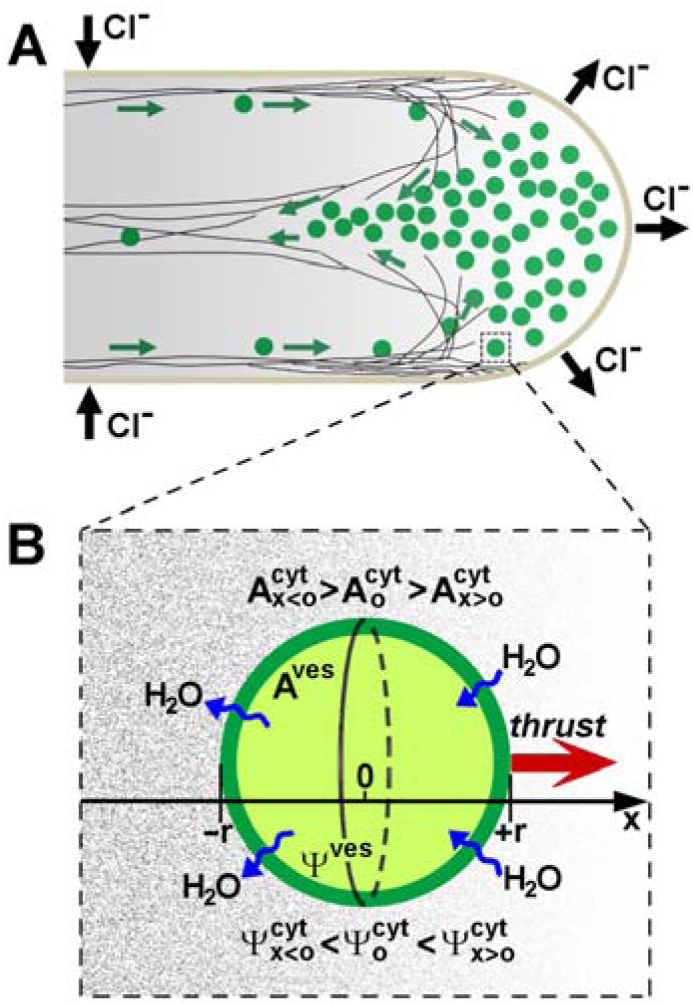
Baseline conceptualization of the effect of transcellular Cl^−^ flux on the vesicle trafficking in pollen tubes (A) Pattern of vesicle motion, arrangement of cytoskeletal arrays, and topographical mapping of Cl^−^ fluxes and gradient (progressive shading) in angiosperm pollen tubes. In the shank region of the tube, vesicles move along well-defined longitudinal cytoskeletal tracks that are crowded in the subapical zone into a cortical fringe. In the apex, cytoskeleton is sparse, implying that vesicle movement is powered by mere hydrodynamic effects. Modified from Chebli et al. (2013), with kind permission from the authors. (B) A vesicle in a solute concentration gradient (progressive shading). In the right half of the vesicle, external solute concentration is reduced, and water enters the vesicle. In the left half of the vesicle, external solute concentration is increased, and water is expelled from the vesicle. Vesicle moves in the direction opposite to the water flow, i.e. toward the pollen tube tip.

Consider a vesicle with a radius *r* located near the flank of the tube apical dome (Fig. 1). Denote osmolarity of the vesicle lumen solution as *A^ves^* and osmolarity of the cytosol surrounding the vesicle as 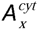, where subscript *x* highlights that, due to the gradient of cytosolic Cl^−^ concentration, the osmolarity of the cytosol varies as a function of longitudinal coordinate *x* (horizontal axis in Fig. 1B). Let us choose an arbitrary origin (*x* = 0) at the tube cross-section that divides the vesicle into equal front and rear halves (right and left halves, respectively, in Fig. 1B). Then, the cytosolic osmolarity at the origin cross-section should be designated as 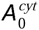. Let us also denote water potential of the vesicle lumen solution as Ψ*^ves^* and water potential of the cytosol as 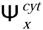, where subscript *x* highlights that the water potential of the cytosol, like its osmolarity, also varies as a function of *x*. If we assume that the vesicle volume is constant, then, as explained in Appendix (2), the following equality holds:

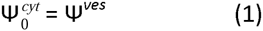

Equation (1) means that water potential of the cytosol near the origin plane (*x* = 0) is equal to the water potential of vesicle lumen solution. In the front half of the vesicle, where cytosolic concentration of Cl^−^ is reduced 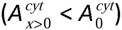, the cytosolic water potential should be higher than the vesicle water potential 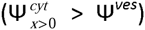, implying a water influx across the front vesicle surface. Similarly, in the rear half of the vesicle, where cytosolic concentration of Cl^−^ is increased 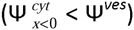, the cytosolic water potential should be lower than the vesicle water potential 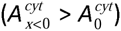, implying a water efflux from the rear vesicle surface. The influx of water through the front vesicle surface and the efflux through the back give rise to the continuous retrograde water flux.

From the above consideration, one can see that the vesicle behaves like a jet engine. When placed in a solute concentration gradient, the vesicle converts the internal energy of the solution into the kinetic energy of the water jet. Momentum conservation requires vesicle propulsion in the direction opposite to the water jet, i.e. toward the pollen tube tip. The conceptual and analytical framework for osmophoretic motion pioneered by Anderson (1983, 1984, see also Anderson 1986, 1989; Nardi et al. 1999) suggests that the vesicle velocity (*v*) is nearly proportional to the vesicle radius (*r*), membrane hydraulic conductivity (*L*), and solute osmolarity gradient 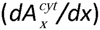:

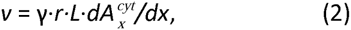

where *γ* is a proportionality constant (Appendix 3). In order to evaluate the proportionality constant, the experimental data of Nardi et al. (1999) can provide a benchmark. The authors observed the movement of artificial DMPC bilayer vesicles of radius 10 μm in a gradient of solute concentration of 10^4^ mol·m^−4^ and found the drift velocity to be 1–2.5 μm per second. According to Orsi et al. (2010), the hydraulic conductivity of DMPC membrane is about 2·10^−13^ m·s^−1^·Pa^−1^ (coefficient of water permeability 14–52 μm·s^−1^). From these data, the proportionality constant *γ* can be estimated as follows:

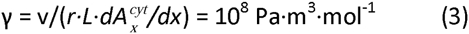

By substituting the calculated constant *γ* and the literature value for hydraulic conductivity of pollen tube plasma membrane (10^−13^ m·s^−1^·Pa^−1^: Sommer et al., 2008; Shachar-Hill et al., 2013) into Equation (2), one can find that a vesicle with a radius of 100 nm would move osmophoretically in a solution with an osmolarity gradient of 10^6^ osmol·m^−4^ at a velocity of about 1 μm·s^−1^.

The above analysis, while being fairly straightforward, is incomplete. It does not incorporate indirect, yet still profound effects that cytosolic Cl^−^ gradient can have on the vesicle dynamics. These effects stem from the fact that cytosolic Cl^−^ gradient creates an osmotic pressure imbalance not only at the vesicle-cytosol interface, but also along the cell plasma membrane and therefore provides a driving force not only for the retrograde water movement throughout the vesicles but also for bulk anterograde transcellular water flow. More specifically, the osmolarity gradient of the order of 3·10^4^ − 2·10^6^ mol·m^−4^ (Appendix 1) implies the difference in osmotic potential between the tube apical cytoplasm and the immediate subapical zone (20 μm apart from the tip) of 2.5 to 100 kPa. This differential should lead to water entrance into the cell across the plasma membrane at the tube shank and water efflux at the cell apex, giving rise to the acropetal transcellular water flow. The rate of this flow can be indirectly estimated based on the literature data for the hydraulic conductivity of pollen tube plasma membrane as 2.5·10^−10^ – 10^−8^ m·s^−1^ (Appendix 4). However, this estimate is likely to be conservative, as the high permeability of the apical plasma membrane for Cl^−^ suggests that its conductivity for water can also exceed the average value of plasma membrane hydraulic conductivity used in the above calculation. This compels one to assume that the water flow can be fast enough to yield a tangible contribution to the vesicle dynamics by means of the advective entrainment of suspended vesicles.

To gain further insight into the effects that cytosolic Cl^−^ gradient has on the vesicle dynamics it is necessary to take into account that the acropetal water flow can bring a plethora of dissolved compounds to the growing cell tip. The majority of these compounds cannot penetrate the apical plasma membrane, so they can accumulate in the apical cytoplasm giving rise to a tip-focused concentration gradient. This pertains primarily to high-molecular compounds whose transport is dominated by bulk hydrodynamic flow rather than diffusion. To quantify this aspect, it is instructive to express the average time *t* required for a molecule with a diffusional coefficient *D* to migrate a distance *x* by using the Einstein–Smoluchowski relation:

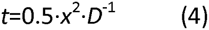

For small ions and molecules (Cl^−^, K^+^, glucose) *D* is of the order of 10^−9^ to 10^−10^ m^2^·s^−1^, and it takes about 1 second for such molecules to diffuse 50 µm. Diffusivity of macromolecules is orders of magnitude lower. The diffusion coefficient for G-actin is estimated to be about 5·10^−12^ m^2^ ·s^−1^ (McGrath et al. 1998; Vitriol et al., 2015), and it takes several minutes for G-actin to diffuse the same 50 μm. This suggests that the diffusion of macromolecules in the retrograde direction cannot offset their anterograde net movement caused by bulk water flow. A possible result is a double-humped profile of cytosolic osmolarity, with the first maximum in the tube shank region, where Cl^−^ enters the cell, and the second maximum near the apical plasma membrane, where high-molecular compounds accumulate. These two zones are separated by an area with lower solute concentration (Fig. 2B).

**Fig. 2.**
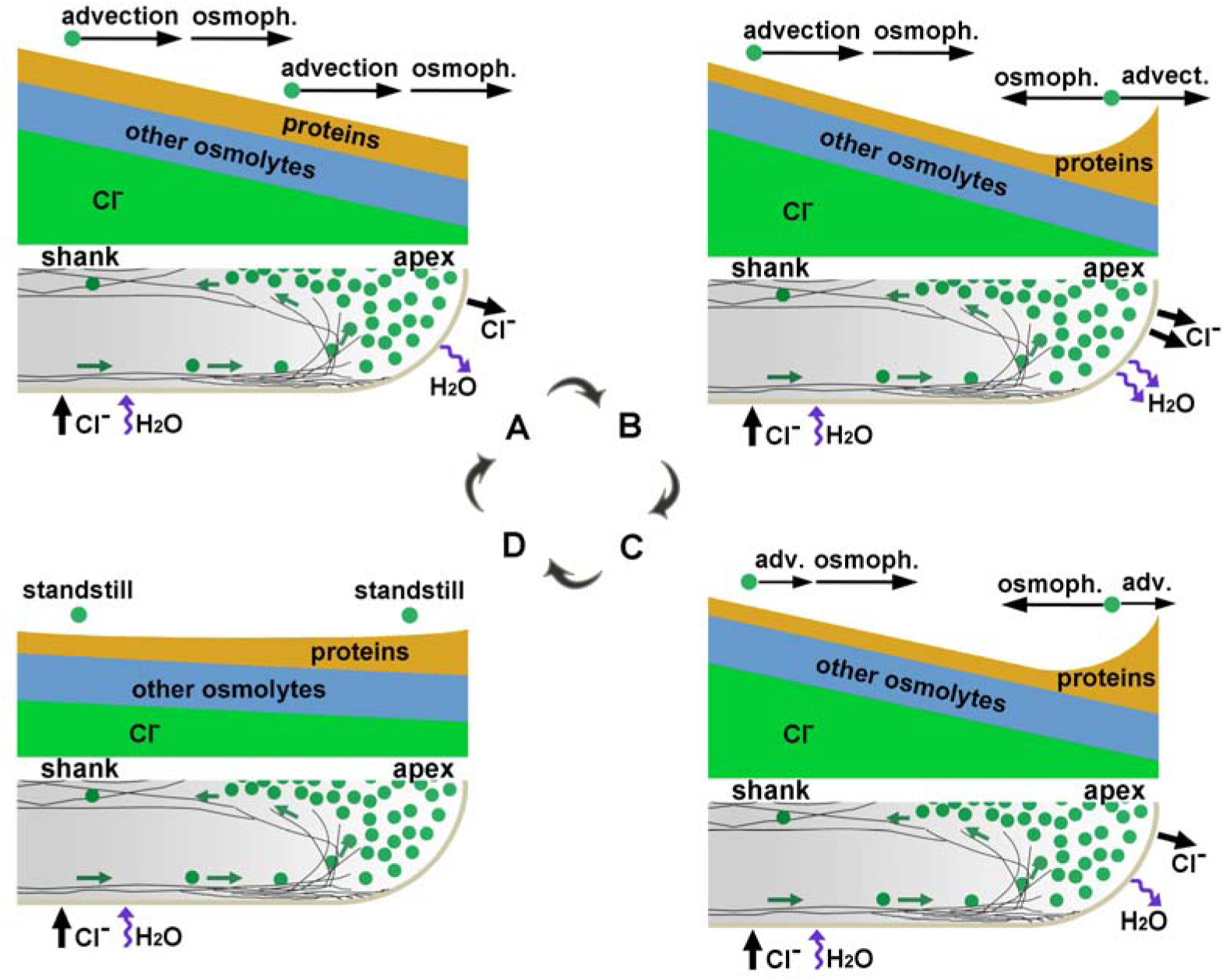
Spatiotemporal variability of cytosolic osmolarity and its impact on the vesicle dynamics. When Cl^−^ and water efflux increases (A), both the bulk water flow and the osmophoresis tend to move the vesicles in the anterograde direction. When Cl^−^ efflux reaches a maximum (B) and decreases (C), the vesicle dynamics in the apical region is dominated by osmophoresis which drives the vesicles in the retrograde direction. When Cl^−^ efflux has ceased (D), there is no hydrodynamic (incl. osmophoretic) forces that could have a tangible effect on the vesicle movement.

The oscillatory character of Cl^−^ efflux implies periodic perturbations in the gradient of cytosolic osmolarity. Because of the high Cl^−^ diffusivity, any variation in apical Cl ^-^efflux would entail immediate changes in the Cl^−^ gradient along the whole pollen tube: more intense efflux, steeper gradient. On the other hand, the low diffusivity of macromolecular osmolytes implies that their gradient varies slowly and depends not so much on the current intensity of the apical water outflow, but on the water flow rate in the immediate past. That is, although it is the spatial profile of Cl^−^ concentration which drives bulk anterograde water flow and eventually causes the accumulation of high-molecular osmolytes in the apical cytoplasm, the Cl^−^ gradient and the gradient of high-molecular compounds can oscillate out of phase. This implies that the gradient of high-molecular osmolytes becomes steep some time after the Cl^−^ efflux has started to steadily increase, reaches the peak value after Cl^−^ efflux has peaked, and persists for a while after rapid Cl^−^ efflux has ceased.

On account of the spatiotemporal variability of the cytosolic osmolarity, it goes to show that osmophoresis can push vesicles either in the anterograde or retrograde direction depending on the vesicle position and current phase of the Cl^−^ efflux cycle. Besides osmophoresis, the water flow is able by itself influence vesicle dynamics and propel their anterograde motion, further complicating the physics involved. Nevertheless, the following qualitative picture may be assumed to have some measure of general validity (Fig. 2). In the tube shank, both the acropetal water flow and the osmophoresis, which here is actuated directly by Cl^−^ gradient, tend to move the vesicles in the anterograde direction. However, since the movement of the vesicles along the tube shank is controlled by motor proteins, the significance of these hydrodynamic effects for the long-distance vesicle transport is unclear. In the cell apex, where cytoskeleton is sparse, the impact of the hydrodynamic effects on the vesicle dynamics should be more profound, albeit also more intricate. It relies on the delicate interplay between, on the one hand, the tip-focused gradient of high-molecular compounds which favors vesicle propulsion in the retrograde direction and, on the other hand, the oppositely directed Cl^−^ gradient and acropetal water flow which both contribute to anterograde vesicle advance. These latter factors generate maximum driving force when Cl^−^ efflux peaks, while the tip-focused gradient of high-molecular compounds produces the most retrograde thrust with a time delay. This phase shift ensures that during each oscillatory cycle anterograde and retrograde vesicle motions can be separated in time.

Besides the changes in the osmolarity gradient, another important factor that can affect the balance between the aforementioned opposing forces is the vesicle radius. According to the Stokes’ law, the drag force (*F_d_*) increases proportionally to the vesicle radius (*r*):

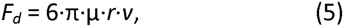

where μ is the dynamic viscosity and *v* is the velocity of the bulk water flow relative to the vesicle. The osmophoretic force (*F_osm_*), in contrast, increases as the third power of the vesicle radius:

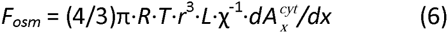

where 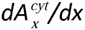 is the osmotic gradient and *χ* is the auxiliary coefficient related to membrane hydraulic conductivity (Appendix 5). By comparing Equations (5) and (6) one can see that the larger the vesicle radius, the greater contribution in the vesicle dynamics makes the osmophoretic force (versus the Stokes’ drag). Since the hydrodynamic drag imparts to the vesicles in the tube apex predominantly anterograde momentum, the power mismatch implies that, under certain conditions, vesicles with smaller radius would move in the anterograde direction while, simultaneously, vesicles with larger radius would move in the retrograde direction. Thereby, large organelles can be strongly repelled from the apex, and this may explain the specific cytology of the tube apical “clear zone”, which is rich in small vesicles but devoid of amyloplasts, vacuoles and other large structures (Helper et al., 2013; Cai et al., 2015; Hepler and Winship, 2015).

The hydrodynamic effects associated with apical Cl^−^ efflux can also have a strong influence on the cytoskeletal dynamics. In particular, the acropetal water flow driven by polar Cl^−^ efflux can carry G-actin from the tube shank to the growing cell tip. In the subapical region of pollen tubes, the actin network is organized into a dense cortical structure referred to as a collar or fringe, where longitudinally oriented microfilaments converge into a palisade, starting 1–5 μm behind the extreme apex and extending basally for additional 5-10 μm (Lovy-Wheeler et al. 2005). The essential contribution of the actin fringe to polarized growth of pollen tubes (Kroeger et al., 2009; Zhang et al., 2010; Bou Daher and Geitmann 2011; Dong et al., 2012; Rounds et al., 2014) suggests that the fringe, during its assembly, may extend apical surface and thereby drive the tip growth. For some cell types, it has also been found that transport of G-actin monomers to barbed ends of actin filaments is the key rate-limiting step in filament elongation and cell motility (Mogilner and Edelstein-Keshet, 2002; Novak et al., 2008; Vitriol et al., 2015). Putting these sets of findings together, it seems likely that the acropetal movement of actin monomers advected by bulk water flow can help explain why oscillatory Cl^−^ efflux and cell growth vary in-phase (Gutermuth et al., 2013).

Another important factor which has to be considered in connection with G-actin transport to the growing tube tip is the electrical polarity of the cell. Breygina et al. (2009b) reported a prominent longitudinal gradient of plasma membrane potential along the growing pollen tubes. The authors provided evidences for a membrane depolarization near the tube tip, and found that anion channel blockers eliminated this longitudinal electrical gradient. These findings are mutually corroborative since the apical Cl^−^ efflux should lead to a local membrane depolarization. Importantly, at physiological pH, G-actin with its isoelectric point of about 5.0 exhibits a net negative charge. Therefore, the apical Cl^−^ efflux, leading to local membrane depolarization, can promote the delivery of negatively charged actin monomers to the growing tube tip. A recent theoretical analysis of the conjunction between electrodiffusion and osmosis with reference to general cell physiology developed by Mori (2012; 2015; Mori et al. 2011) may provide a promising mathematical framework for capturing the interrelated impact of electrophoresis, diffusion and osmosis on ion movements.

The massive tip-localized efflux of osmolytes is not peculiar to pollen tubes, but is also a feature of other tip-growing cells. Over 40 years ago, Nuccitelli and Jaffe (1974) observed strong tip-focused ionic currents in growing rhizoids of fucoid zygotes. Subsequent experiments demonstrated that these ionic currents involved high-capacity efflux of Cl^−^ and K^+^ at the growing rhizoidal tip, with Cl^−^ efflux dominated and occurred in a pulsatile manner (Nuccitelli and Jaffe, 1976a). The average magnitude of Cl^−^ efflux depended on cell turgor pressure and was sometimes as high as 50 pmol·cm^−2^ ·s^−1^ (Nuccitelli and Jaffe, 1976b). Although this value is 3 orders of magnitude below the peak magnitude of Cl^−^ efflux from pollen tube tips (Zonia et al., 2002; Tavares et al., 2011), such a comparison requires great caution due to differences in the experimental design and methods employed. It should also be taken into account that fucoid rhizoids have growth rates that are typically 10-to 100-fold less than growth rates of pollen tubes. It therefore seems plausible that, in the former case, weaker ion fluxes and lower osmotic gradients can ensure adequate forces for advection of growth-limiting proteins (such as G-actin) and propulsion of vesicles to the growing rhizoidal tips.

The above considerations suggest that in tip-growing cells, osmotically generated forces make a significant contribution to vesicle and cytoskeletal dynamics, thereby promoting cell motility and polarized growth. A corollary is that tip-growing cells can dodge the extension-induced structural instability by targeting vesicle delivery strategically to the weakest domains of the nascent cell wall. Indeed, the membrane stretch is able to activate not only bona fide mechanosensitive ion channels but also a variety of voltage-, ligand- and Ca^2+^-gated channels (Anishkin et al., 2014) and therefore can directly or indirectly focus Cl^−^ efflux to the tenuous cell wall regions. The polarized Cl^−^ efflux can navigate vesicle trafficking accordingly, leading to targeting of cell wall constructional materials to the most stressed and loosened domains at the cell surface. Noteworthy, it was found that Cl^−^ efflux from rhizoidal tips can be strongly stimulated by membrane stretch (Nuccitelli and Jaffe 1976b). At the same time, the vesicle motility in the growing rhizoidal tips is also highly sensitive to the plasma membrane stretch (Brownlee et al., 1998). Taken together, these data suggest that the osmophoresis provides a neat way to couple the intensity and direction of vesicle trafficking with cell wall mechanical status, with a possible role in prevention of turgor-triggered cell bursting.

## Appendix 1

Fick's law relates the diffusive flux density *J* to the concentration gradient *dC*/*dx* and the diffusion coefficient *D* as:

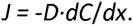

Assuming that the diffusion coefficient for Cl^−^ in cytosol is about 10^−9^ m^2^·s^−1^ (2 times smaller than in water: Kushmerick and Podolsky, 1969) and substituting this coefficient and the peak magnitude of oscillatory Cl^−^ efflux (1 - 60 nmol·cm^−2^ ·s^−1^) into above equation gives the concentration gradient near the pollen tube apical dome ranging from 10^4^ to 6·10^5^ mol·m^−4^. The continuity and uniformity of the ion current between the flanks of the apical dome and the shank domain where Cl^−^ enters the cell requires that near the flanks of the apical dome and behind the concentration gradient exceeds the gradient near the apical dome by a geometrical factor *G*:

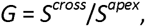

where *S^cross^* is the area of transverse section across the flanks of the apical dome, and *S^opex^* is the dome surface through which the efflux occurs. For a prolate hemispheroid apex the factor *G* can be estimated as 3, implying the concentration gradient from 3·10^4^ to 2·10^6^ mol·m^−4^ in the region spanning from the flanks of the apical dome to the beginning of the shank zone where Cl^−^ influx is started.

## Appendix 2

Let us assume that the vesicle volume is constant. Then the total molar water flux *J_Σ_* (units: mol·s^−1^) across the vesicle surface is zero:

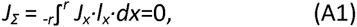

where subscript *x* highlights that *J_x_* (units: mol·s^−1^ ·m^−2^) is the molar flux density of water through the circumference of length *l_x_* at the vesicle surface with a longitudinal coordinate *x* (Fig. 1B).

*J_x_* can be expressed as:

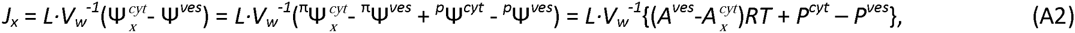

where *L* is the membrane hydraulic conductivity, *V_w_* is the molar volume of water (18·10^−6^ m^3^ ·mol^−1^), 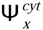and Ψ*^ves^* are water potentials of the cytosol (in a point with longitudinal coordinate *x*) and of the vesicle solution, respectively, 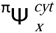 and *^π^*Ψ*^ves^* are respective osmotic potentials, 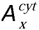 and 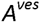 are respective solute osmolarities, *^p^*Ψ*^ves^* and *^p^*Ψ*^ves^* are respective pressure potentials, *P^cyt^* and *P^ves^* are respective hydrostatic pressures, *R* is the gas constant, and *T* is the absolute temperature.

Substituting the local water flux *J_X_* from Equation (A2) into Equation (A1) gives:

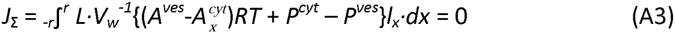

A linear gradient approximation can be applied to describe the dependence of the cytosolic osmolarity on the longitudinal coordinate *x* as:

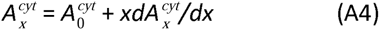

The expression for cytosolic osmolarity from Equation (A4) can be inserted into Equation (A3) to give:

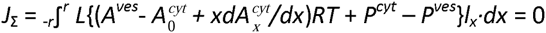

The above integral can be split as a sum of three integrals:

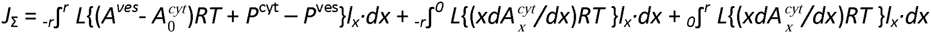

The limits of integration in the right-hand integral can be reversed with a corresponding sign change:

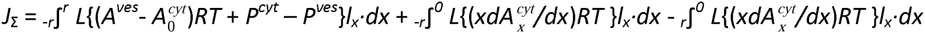

From geometrical considerations (Fig. 1B) one can see that the second and third integrals are equal, and therefore Equation (A4) can be reduced to:

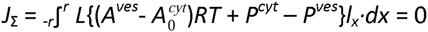

This equation can be solved as:

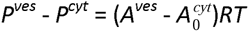

The last equation can be rewritten in terms of water potential:

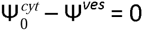

## Appendix 3

Anderson (1983) derived the following expression for the velocity *v* of a semipermeable vesicle of radius *r* moving in the solute osmolarity gradient *dA^out^* /d*x*:

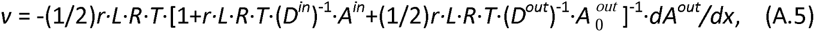

where *D^in^* and *D^out^* are the solute diffusion coefficients inside and outside the vesicle, respectively. Substitution of the pertinent values for *L* (10^−13^ m·Pa^−1^ ·s^−1^), *R* (8.31 m^3^·Pa·K^−1^·mol^−1^), *T* (300 K) and *D* (10^−9^ m^2^ ·s^−1^) in Equation (A.5) shows that for endosomal vesicles the second and third summands are negligible. Therefore, in the present context, Equation (A.5) can be reduced to:

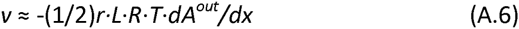

Independently of Anderson (1983), Nardi et al. (1999) suggested the following expression for the osmophoretic vesicle velocity:

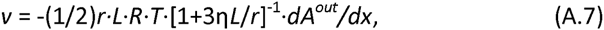

where η is the solvent viscosity. Substituting the relevant values for *L*, *r* and η shows that 3η*L*/*r* << 1 and Equation (A.7), just as Equation (A.5), can be reduced to Equation (A.6).

The experiments of Nardi et al. (1999) revealed that Equation (A.7) greatly underestimates the efficiency with which vesicles are able to transform osmotic gradients into mechanical motion. The reason of such a discrepancy deserves detailed analysis, which is, however, complicated by the lack of acceptable kinetic interpretation of osmosis. An operational way to overcome this complexity is to assign an empirical factor γ calculated based on the experimental data of Nardi et al. (1999) as follows: 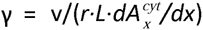.

## Appendix 4

The volume flux density of water across the plasma membrane in the cell apex 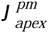 (units: m·s^−1^) can be expressed as:

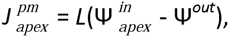

where *L* is the membrane hydraulic conductivity, 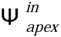 and Ψ*^out^* are water potentials of the apical cytosol and of the outer milieu, respectively. The area of the apical plasma membrane (where Cl^−^ and water exit the cell) is smaller than the plasma membrane area in the tube shank (where Cl^−^ and water enter the cell), and therefore 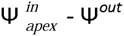 appears to be higher than 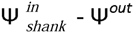. This suggests that, at a first approximation, Ψ*^out^* can be replaced by 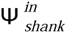, yielding:

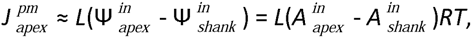

Where 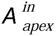 denotes apical cytosolic osmolarity and 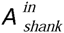 denotes the osmolarity in the subapical zone, 20 μm apart from the tip. The above values for Cl^−^ gradient of 3·10^4^ to 2·10^6^ mol·m^−4^ (Appendix 1) imply the differential between 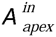 and 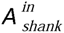 to be about 1 – 40 мМ. Substituting in the above equation this differential and the hydraulic conductivity of pollen tube plasma membrane (10^−13^ m·s^−1^ ·Pa^−1^: Sommer et al., 2008; Shachar-Hill et al., 2013) gives the water flow rate of the order of 2.5·10^−10^ – 10^−8^ m·s^−1^.

## Appendix 5

Osmophoretic force (*F_osm_*) refers to the force equal and opposite to the one that has to be applied to a vesicle to prevent its osmotically driven movement. An analytical expression for this force can be derived in two steps. The first step is to calculate the maximum thermodynamically permitted force acting on a semipermeable vesicle in a solute concentration gradient by assuming that the transmembrane osmotic pressure difference exerts equal mechanical pressure which causes the vesicle to move. The second step is to refine the derived equation by compensating the error incurred by the premised assumption.

The maximum thermodynamically permitted osmophoretic force can be expressed as the function of the vesicle radius and osmolarity gradient as follows:

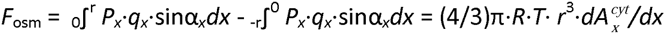

where *P_x_* is the pressure in a point with a longitudinal coordinate *x*, *q_x_* is the perimeter of the circle at the vesicle surface with a coordinate *x*, *α_x_* is the angle between the *x*-axis and the vesicle surface in a point with a coordinate *x*, *R* is the gas constant, *T* is the absolute temperature, *r* is the vesicle radius, and 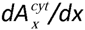 is the osmolarity gradient. The bias incurred by assuming the equality between transmembrane osmotic and mechanical pressures can be corrected by introducing into above equation the factor *L*·*χ*^−1^ which is the ratio between actual membrane hydraulic conductivity and the hydraulic conductivity in an ideal case when water flow is restricted by inertial forces but not molecular friction. The result is:

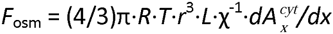

